# Metal-catalyzed phosphorylation by phosphite at the origin of bioenergetics

**DOI:** 10.64898/2026.05.13.724781

**Authors:** Nadja K. Hoffmann, Manon L. Schlikker, William F. Martin

**Author notes:** These authors had equal contribution (Nadja K. Hoffmann, Manon L. Schlikker).

## Abstract

How did phosphate become the universal energetic currency of life? Traditional approaches to phosphorylation in early evolution studies entail oven drying, non-aqueous solvents, dangerously reactive forms of phosphorus, or other non-physiological conditions. With microbial physiology as a *vade mecum*, we have recently found that phosphite, HPO_3_^2–^, which is enzymatically oxidized by many microbes and which naturally occurs in serpentinizing hydrothermal vents, will readily phosphorylate ribose, glucose, glycerol, serine, AMP, creatine and acetate to generate phosphoester, phosphoanhydride and acylphosphate bonds in hours to days at 25-100°C in pure alkaline water. These reactions are thermodynamically favourable because anoxic phosphite oxidation to phosphate and H_2_ is highly exergonic, but they do not proceed without catalysts. The most effective catalyst yet identified is a nanoparticular form of a shiny metal: zero-valent (native, or elemental) palladium (Pd^0^). Native palladium, like phosphite, also naturally occurs in serpentinizing hydrothermal vents, as do other native platinum group elements (PGE), including Pt, Rh, Ru and Ir. Here we test those PGE as catalysts of phosphite oxidation and phosphorylation. Though all metals tested readily oxidize phosphite, only Pd^0^ efficiently catalyzes phosphorylation, generating phosphorylated products at concentrations often equal to their physiological concentrations in growing *Escherichia coli* cells. Metaphosphate is a possible reaction intermediate. In phosphorylation reactions via phosphite oxidation (D*G*_0_′= –46 kJ·mol^−1^), a portion of the energy released is conserved in phosphorylated products, as in biological energy conservation. A natural environment and energy-conserving thermodynamics implicate these facile aqueous phosphorylating reactions in the origin of bioenergetics.

## Introduction

Bioenergetics and genetics are both built on phosphate [1], making phosphate indispensable at origins. In modern life, phosphate has a structural role as an integral component of RNA, DNA and phospholipids, with phosphorus comprising about 2%of the dry weight of an *E. coli* cell [2]. It also has a functional role, serving as an energy currency, mainly in ATP, the hydrolysis of which to ADP and P_i_ is exergonic with D*G*_0_′–32 kJ·mol^−1^, such that ATP hydrolysis can be enzymatically coupled to otherwise endergonic reactions so as to render the overall reaction thermodynamically favourable [3]. In some reactions, ATP is hydrolyzed to AMP and pyrophosphate, which is rapidly hydrolyzed by pyrophosphatases, rendering reactions irreversible under physiological conditions [4,5]. In ‘genetics first’theories of origins, the role of phosphate as a component of nucleic acids usually stands in the foreground. In ‘metabolism first’theories of origins, the role of phosphate is seen in energy conservation and allowing uphill reactions to go forward. The relative contributions significance of phosphate’s functions in energy metabolism and structure during the growth process can be quantified. During cell division, *E. coli* hydrolyses ∼10^10^ phosphate bonds in ATP to make a cell that contains ∼10^8^ phosphate residues (∼20 million in phospholipids and ∼80 million in RNA) [6], revealing a 100:1 predominance of phosphate’s use in bioenergetics over use in biomolecular structure. That ratio is probably as old as life itself. How did phosphate come to assume its central role in bioenergetics?

Obviously, routes to incorporation of phosphate into organics had to proceed the existence of RNA, it cannot be any other way. This has placed a premium on the identification of abiotic chemical routes that can lead to the synthesis of organophosphates, in particular ribonucleoside phosphates, under prebiotic conditions [7]. That chemistry has proven to be anything but simple. Traditional routes of non-enzymatic, prebiotic-type phosphorylation include the incubation of phosphate in pure formamide [8], the use of non-physiological activating reagents such as cyanoacetylene, cyanogen, cyanamide, cyanoformamide, or cyanate [9], oven-heating with phosphate to dryness with or without urea [10–12], or the use of highly reactive forms of phosphorus as schreibersite, (FeNi)_3_P, a phosphide that occurs in meteorites [13], or the magma-based synthesis of P_2_O_5_ [14], which reacts violently in contact with water. These approaches do not converge on a single kind of chemistry or environment as a source of phosphate at origins.

Our approach to phosphorylation in early chemical evolution stems from our interest in (i) acetogenesis and methanogenesis as the most primitive forms of microbial energy metabolism and (ii) serpentinizing hydrothermal vents as a possible site for the origin of metabolism and life [15–18]. Serpentinization is a geochemical process that entails circulation of deep-sea water through the Earth’s crust and chemical reactions of water with Fe(II) rich minerals (rock water-interactions) that generate highly reducing conditions (pH 9-11) and H_2_ gas as diffusible reductant [19–22]. The highly reducing conditions of serpentinizing hydrothermal vents lead to the formation and deposition of shiny transition metals (syn. native metals, zero-valent metals, elemental metals), mainly Ni^0^, Fe^0^, Co^0^, Ni_3_Fe alloys, CoFe alloys [23], but including the less common platinum group elements (PGE), Pd^0^, Pt^0^, Rh^0^, Ru^0^, Ir^0^ and Os [24]. Previous work had shown that Ni^0^, Co^0^, Fe^0^ and their alloys serve as effective catalysts for H_2_ dependent CO_2_ fixation in reactions that specifically generate products of the acetyl-CoA pathway [25]: formate, acetate, pyruvate and methane under aqueous conditions of serpentinizing vents [26-30]. The reactions generate up to 200 mM formate and up to 200 µM pyruvate [27], equal to the physiological concentration of pyruvate in cells that use that acetyl-CoA pathway for carbon and energy metabolism [31]. Key enzymes of the acetyl-CoA pathway use Fe, Ni and Co catalysis at their active sites [32–35], but the catalysis of the enzymes can be replaced by metallic state Fe, Co and Ni themselves [17]. Native metals reduce NAD^+^ [36, 37], ferredoxin [38] and reductively aminate both pyridoxamine [39] and seven different 2-oxoacids to the corresponding amino acids [40] in addition to catalyzing the reactions of the reverse TCA cycle [40, 41]. These native metals replace not just enzymes, they also replace cofactors [42]. We have found Ni^0^ to be conspicuously active as a broad-specificity catalyst of metabolic reactions [43].

Because native transition metals can replace enzymes and cofactors in ancient biochemical pathways, we asked whether they could also replace the enzyme AdpA, which catalyzes a unique, phosphite-dependent NADH-forming substrate level phosphorylation reaction:

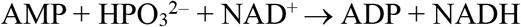

which is the sole source of phosphate in the bacterium *Phosphitispora fastidiosa*, a bacterium that uses the acetyl CoA pathway [44]. The AdpA reaction is all the more notable in that it represents *a natural route of phosphate incorporation* (*sensu* the origins problem), but in a reaction that does not use phosphate as a substrate, using phosphite instead and converting phosphite to phosphate in a redox reaction during the enzymatic mechanism [44]. The use of phosphite by AdpA also directly connects this reaction to serpentinizing hydrothermal vents, because (i) serpentinized rocks can contain substantial proportions of phosphite, in addition to phosphate [45] and (ii) because phosphite-oxidizing genes are enriched in communities that inhabit serpentinizing hydrothermal vents [18, 46, 47]. We found that Ni^0^, an effective catalyst of redox reactions, catalyzes the oxidation of phosphite to phosphate and replaces the enzyme in the AdpA reaction, also replacing NADH with H_2_ [42, 48]. But we also found that Pd^0^, directly below Ni^0^ in the periodic table, is a far more effective catalyst of the same reaction:

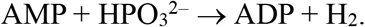

The formation of the phosphoanhydride bond in ADP in that reaction is energetically driven by the highly exergonic phosphite oxidation reaction:

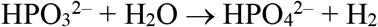

with D*G*_0_′= –46 kJ·mol^−1^ [49]. Phosphite of Pd^0^ phosphorylates AMP, but it also phosphorylates serine, ribose, glucose, cytidine, glycerol, creatine and acetate [48] as shown in **Figure 1**.

**Figure 1.**
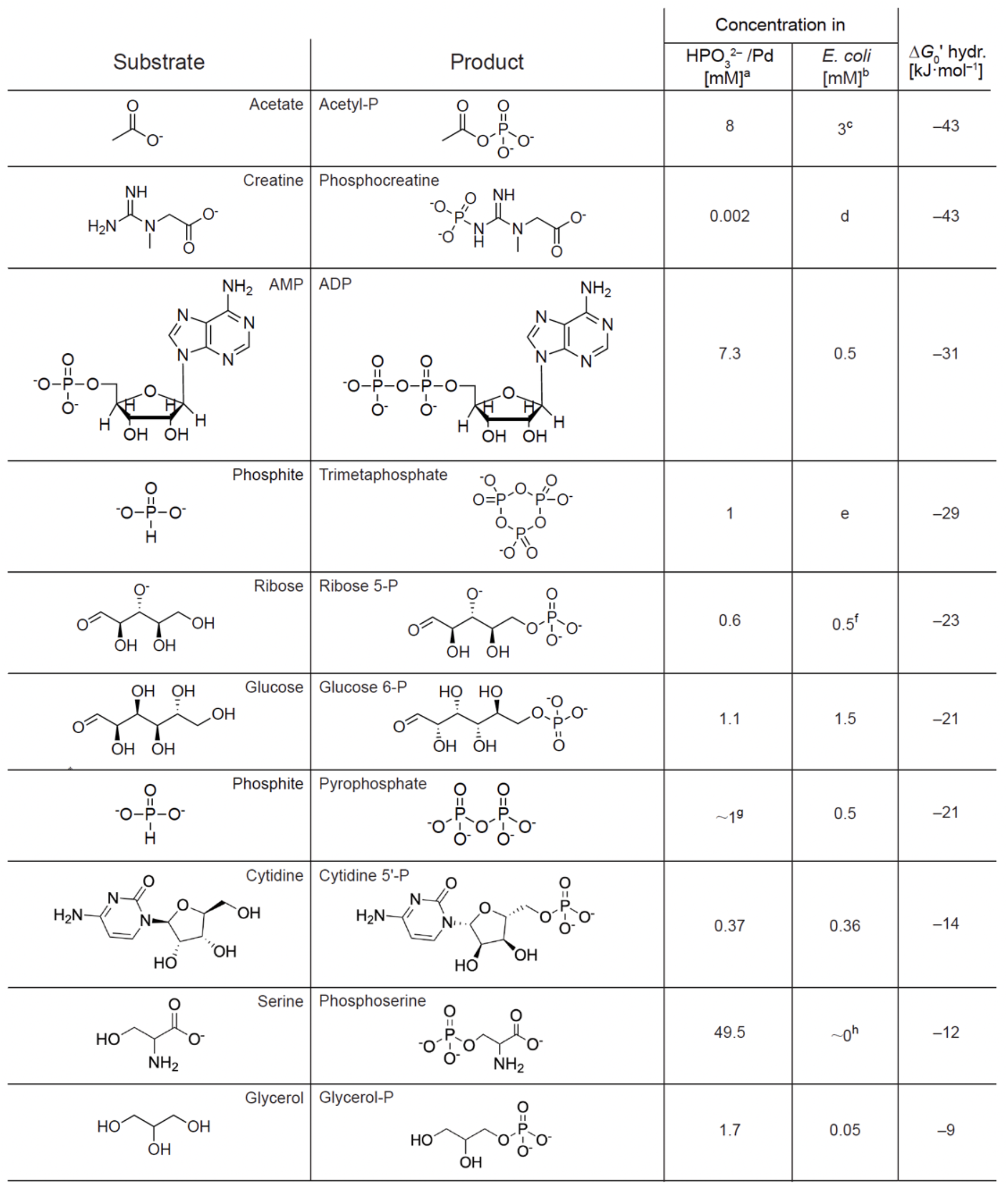
Products obtained by Schlikker *et al*. (2026) [48] in reactions of each substrate (100 mM) with 200 mM phosphite in water over Pd^0^ catalysts. Times 18–72h, temperatures 25–80°C (details in **Schlikker** *et al*. (2026) [48]). ^a^) Concentrations of the phosphorylated product obtained with HPO_3_^2–^ over Pd^0. b^) Cytosolic concentration of the product in *E. coli*, values from **Bennett** *et al*. (2007) [56] unless otherwise indicated. ^c^) From ref. **Klein** *et al*. (2007) [57]. ^d^) Does not occur in *E. coli*, a typical energy storage compound (phosphagen) of animal cells [58];concentration determined by LC-MS [48]. ^e)^ Meyerhoff *et al*. (1953) [59] measured the free energy of hydrolysis of trimetaphosphate to three orthophosphate molecules as –88 kJ·mol^−1^ (–29.3 kJ·mol^−1^ per metaphosphate bond). ^f)^ Estimated, **Bennett** *et al*. (2007) [56] report 1.3 mM for ribose-5-P, ribulose-5-P and xylulose-5-P combined. ^g)^ Small specific peak in ^31^P NMR above detection limit of 1%conversion [48];*E. coli* value from ref. [60]. *E. coli* synthesizes cysteine via *O*-acetyl serine and expresses phosphoserine phosphatase that keeps phosphoserine levels low;archaea synthesize cysteine via *O*-phosphoserine but ligated to tRNA [61].

### Are phosphorylated products obtained in relevant amounts?

Phosphite has long been discussed as a theoretical source of phosphate at origins [50–52], but the discussion became more concrete with the isolation of bacteria that use phosphite as their sole source of phosphorus and electrons [53], which prompted the suggestion by **Buckel** (2001) [54] that phosphite oxidation could readily give rise to acetyl phosphate, an ancient currency of biochemical energy [15, 55]. Subsequent studies reported evidence for phosphite in Hadean oceans [13] and detected phosphite in modern serpentinizing systems [45]. The discussion of phosphite as a source of phosphate and phosphorylation, at origins became even more concrete with the recent demonstration that Pd^0^ catalyzes phosphite-dependent phosphorylation [48].

Though Pd^0^ phosphorylates a broad spectrum of substrates in water at mild temperatures, the yields in most cases were low, in the mM range (**Figure 1**). Low yields might lead to questions about their impact at origins. Notwithstanding the circumstance that in a serpentinizing environment phosphite supply would be constant, rather than intermittent (as in meteoritic delivery), from a ‘metabolism first’perspective, the question is whether the yields are *physiologically* relevant, which is easy to check, at least for *E. coli* [62]. In **Figure 1**, we have indicated the concentration of the corresponding product measured from the cytosol of growing *E. coli* cells next to the concentration of phosphorylated product obtained with HPO_3_^2–^ in water.

In most cases, the concentrations of phosphorylated products obtained with HPO_3_^2–^/Pd correspond surprisingly well with the physiological concentrations of the phosphorylated metabolites in *E. coli* (**Figure 1**). We are not suggesting (yet) that metabolite concentrations in *E. coli* are a relic of primordial phosphorylation reactions. Rather, we wish (i) to underscore the (known) observation that concentrations of metabolites involved in core carbon and energy pathways in growing cells of *E. coli* and other microbes are typically in the low mM to low µM range [56] and (ii) to contrast that observation to the new and surprising finding that a single native metal catalyst (Pd^0^) that naturally occurs in serpentinizing hydrothermal systems [24] generates, on the whole, very similar concentrations of phosphorylated products, but not using phosphate or phosphoryl donors as a reactant, rather using HPO_3_^2–^, which also occurs in serpentinized systems [45].

The correspondences of concentrations in **Figure 1** might be coincidental and it should be stated that the reactions shown in the figure used 200 mM phosphite as a reactant although 75 mM HPO_3_^2–^ also works in some cases [48]. But this should not distract from the observation that, in contrast to traditional prebiotic phosphorylation methods, HPO_3_^2–^/Pd^0^ reactions reported so far operate under what one might call ‘generally physiological prebiotic conditions’: water, pH 9, without nitriles or other condensing agents, in hours or days, at temperatures of 20–80°C and without recourse to drying cycles. The latter point is important, because one well established school of thought on origins [63, 64] infers the existence of evaporation based wet-dry cycles on the early Earth to promote phosphorylation for RNA synthesis, based on laboratory syntheses. That is one view. As we see it, whatever prebiotic, environmental phosphorylation mechanism operated at the onset of metabolism and genetics, it had to *operate continuously* until enzymes arose that could usurp the function of the prebiotic phosphorylation mechanism. If drying was required at every phosphorylation step [63, 64], active enzymes never could have arisen, because they denature upon heat drying—recall that ribosomal protein synthesis requires 4 ATP per peptide bond. Phosphorylation under conditions of serpentinization shares several properties with the aqueous chemistry of cells that traditional phosphorylation protocols lack.

An additional observation emerges from **Figure 1**. Traditional approaches to phosphorylation start from phosphate with the goal of forming bonds between phosphate and hydroxyl, phosphoryl, or carboxylate residues on substrates. In doing so, such procedures must add energy in some form so as to overcome the free energy of hydrolysis shown in the right hand column of **Figure 1**. Phosphorylation through phosphite oxidation over metals forges organophosphate bonds in exergonic reactions that conserve a portion of the energy released as phosphoester, phosphoanhydride or acylphosphate bonds, exactly as in metabolism, but using a single shiny (native) metal as the catalyst instead of different substrate-specific enzymes. The highly exergonic nature of the phosphite oxidation reaction and the broad specificity of the native metal catalyst are desirable attributes for a prebiotic energy conserving phosphorylation reaction [7,51], as are the natural co-occurrence of phosphorylating agent and activating catalyst in the same aqueous environment [42, 48]. Perhaps nature did not choose phosphate [1], perhaps there were no viable alternatives in the environment where metabolism and life arose.

### Do other PGEs also catalyze phosphorylation with HPO_3_^2–^?

In the vast literature on phosphate at origins, catalysts have played a very minor role. In biology, phosphite requires enzymatically catalyzed oxidation for the formation of phosphoanhydride bonds [44]. In organic chemistry, PGE are renowned for their catalytic properties, whereby most uses of PGE are in homogeneous catalysis: one atom of the metal is coordinated individually as a charged cation within a soluble organic molecule, sometimes more than one metal atom per molecule [65], pincer complexes being one example catalyst class [66], metal-organic frameworks [67] being another. The metals that we are using as catalysts are nanoparticles, either on supports such as carbon or aluminum silicate or as pure metal nanoparticles. A typical 2 nm diameter Pd nanoparticle will have a few hundred atoms, most commercial nanoparticles are larger. Importantly, the Pd particles we use are not dissolved, they are in the solid phase, they have a concentration of zero in solution and their structure is amorphic (unknown), so that the number of catalytically active atoms per particle, if any, is obscure. Since Pd is an effective phosphorylating catalyst, a brief look at other PGE is worthwhile, because they are deposited and enriched in the same serpentinizing hydrothermal systems as Pd, often as very small particles [24]. Therefore, we investigated Pt, Ru, Rh and Ir in comparison to Pd for their ability to oxidize phosphite and catalyze two phosphorylation reactions: AMP ®ADP and serine ®phosphoserine.

## Material and Methods

Reactions contained 200 mM sodium phosphite (NaH_2_PO_3_;Merck, Sigma-Aldrich) dissolved in distilled water and adjusted to a final volume of 1.5 mL. Where indicated, serine (100 mM;Carl Roth) or adenosine monophosphate (AMP, 100 mM;Merck, Sigma-Aldrich) was added. Catalysts were purchased from Merck, Sigma-Aldrich, unless stated otherwise. Metals (Pt, Pt/Al_2_O_3_, Ir, Rh, Rh/Al_2_O_3_, Ru, Ru/C, Ru/Al_2_O_3_, Pd, Pd/C, Ni and Ni/C) were weighed out under ambient laboratory conditions and added at a loading of 1.5 mmol metal per reaction. Supported catalysts, Ru/C (10 wt%Ru on activated carbon) and Pd/C (10 wt%Pd on activated carbon), were added at amounts corresponding to approximately 0.15 mmol of elemental metal per reaction. Catalyst loadings are reported as mmol of elemental metal. Reaction mixtures were transferred into 3 mL glass vials, sealed with screw caps, and punctured with a sterile needle to allow pressure equilibration while maintaining a closed reaction environment. The vials were placed in a stainless-steel pressure reactor equipped with a temperature controller (Berghof BR-300 with BTC-3000). The reactor was purged three times with argon and subsequently pressurized to 5 bar Ar (99.999%). Reactions were conducted at 100 °C for 72 h. After completion, the reactor was depressurized, and the vials were immediately transferred to 2 mL microcentrifuge tubes. Samples were centrifuged for 20 min at 16,060 ×g to separate the catalyst from the supernatant. The supernatants were collected for NMR analysis.

## Results

To assess whether phosphite oxidation and phosphorylation activity are general properties of transition metal catalysts or restricted to specific metals, a comparative screen of platinum group element (PGE) and nickel-based catalysts was performed. As shown in **Figure 2**, all platinum group element (PGE) catalysts tested readily promoted phosphite oxidation under hydrothermal conditions, whereas only selected metals catalyzed detectable phosphorylation reactions.

**Figure 2.**
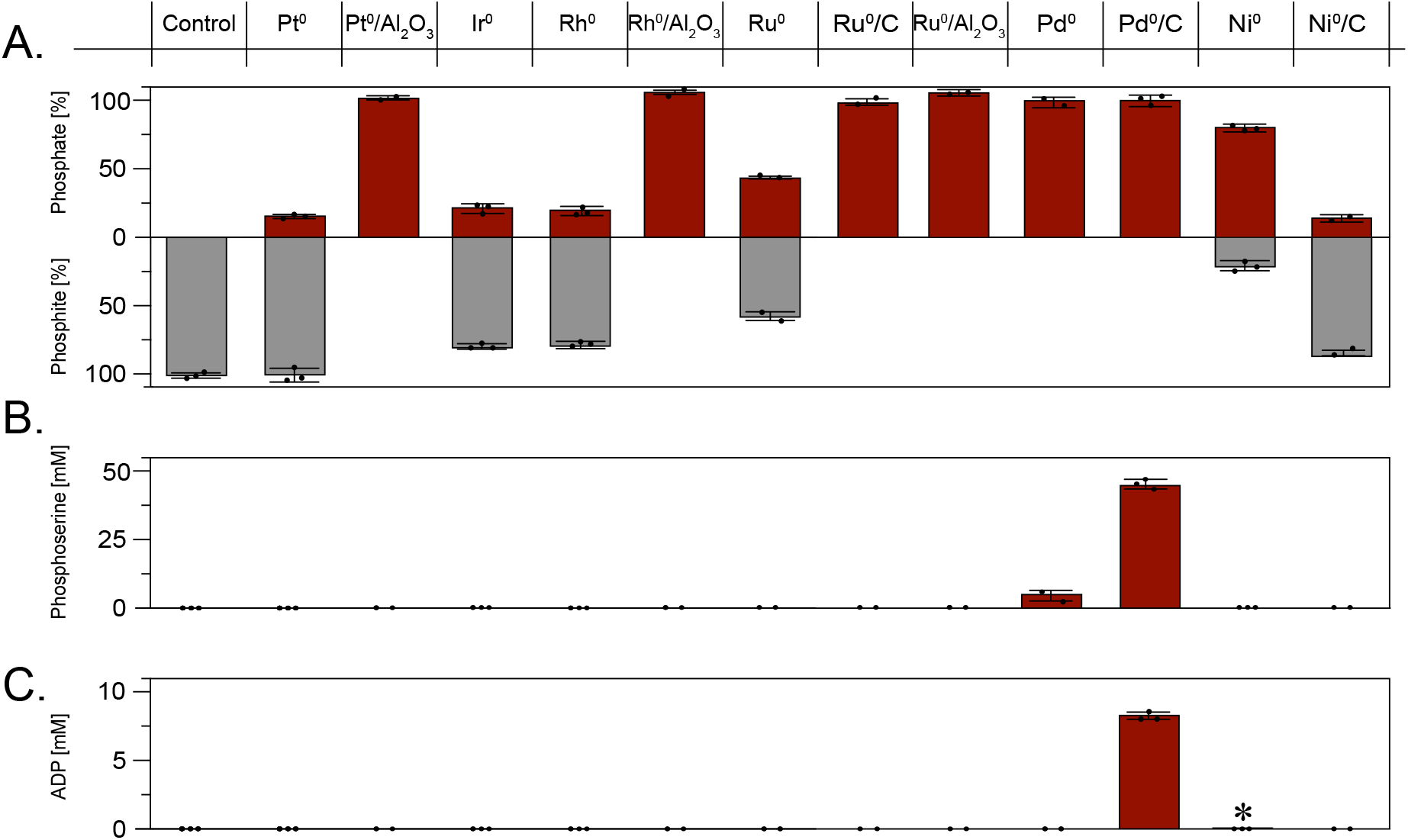
Comparison of platinum-group-element (PGE) and nickel-based catalysts for phosphite oxidation and phosphorylation reactions, control without catalyst at left. **A**, Phosphite oxidation to phosphate in the presence of different metal catalysts after incubation at 100 °C for 72 h under 5 bar Ar in 1.5 mL aqueous solutions containing 200 mM sodium phosphite (^31^P-NMR raw spectra shown in **Fig. S1**). **B**, Formation of phosphoserine from serine and phosphite after incubation at 50 °C for 18 h (^31^P-NMR raw spectra shown in **Fig. S2**). **C**, Formation of ADP from AMP and phosphite after incubation at 50 °C for 18 h (^31^P-NMR raw spectra shown in **Fig. S3**). All reactions contained 200 mM sodium phosphite. Supported catalysts were applied at loadings corresponding to approximately 0.15 mmol elemental metal, except for Pd/C, which was applied at 1.5 mmol. The asterisk (*) indicates the formation of approximately 7 µM ADP, previously detected by LC–MS analysis in Schlikker *et al*. (2026) [48], whereas all other product concentrations were quantified by NMR spectroscopy. Osmium was not investigated owing to laboratory safety considerations.

All metals tested readily oxidized phosphite, probably through hydride transfer coupled to H2 formation (see proposed mechanism in the following section). However, only Pd^0^ efficiently catalyzed the formation of phosphorylated products, including ADP and *O*-phosphoserine, as detected by ^31^P NMR spectroscopy. Because ^31^P NMR has a relatively low sensitivity corresponding to approximately 1%conversion, lower levels of phosphorylation activity cannot be excluded. Nevertheless, this initial low-sensitivity exploratory survey with two substrates suggests that phosphorylating activity may be specific to Pd among the platinum group elements tested. This behavior may relate to the unique electronic and hydrogen-binding properties of Pd, with its filled 4*d*^10^ shell and its strong tendency to form surface and bulk hydride species, which may facilitate phosphorus transfer chemistry under our reaction conditions. Ni^0^ (3*d*^8^4*s*^2^) exhibited minor phosphorylating activity, representing an intermediate behavior between Pd and Pt under the conditions tested.

The carbon support itself is not responsible for phosphorylation, as neither Ru^0^/C or Ni^0^/C generated detectable phosphorylated organics. Also, there is ten times less metal in the reaction when we use carbon supported metals, the C support environment is apparently conducive to phosphorylation in the case of Pd^0^, other supports need to be tested. Phosphite oxidation is highly exergonic (–46 kJ per mol of phosphate produced), hence the equilibrium should lie far on the side of phosphate. In the presence of several catalysts tested, the reaction goes to completion, but for several catalysts, over 50%of the phosphite did not react (**Figure 2A**). This suggests either very slow reaction rates or the presence of reactions that poison the catalyst surface. In the absence of catalysts, phosphite does not undergo oxidation to phosphate in water under anaerobic conditions. The aluminum silicate supported metals are effective catalysts for phosphite oxidation to phosphate, but do not yield phosphorylated products detectable by ^31^P-NMR (**Figure 2**). As noted above, ^31^P-NMR is not a sensitive method, it will only detect phosphorylated products at about 1%conversion rate (2 mM yield in the present case). In the case of Ni^0^, liquid chromatography and mass spectrometry (LCMS) detected 6.7 µM ADP product (0.0067%conversion) (Mrnjavac et al. 2026 [42]), indicated by an asterisk in **Figure 2C**. It is possible that other catalysts screened here will reveal phosphorylated products with LCMS detection. Abiotic-type reactions should ideally be robust in the sense of delivering yields that are relevant for microbial metabolism (**Figure 1**).

## Discussion

### A possible mechanism

Solid state catalysts can be very effective in promoting reactions, the mechanisms of which typically remain very difficult to probe with standard laboratory tools. A glaring exception is Ertl’s elucidation of N_2_ fixation and CO oxidation mechanisms involving direct observations of reaction intermediates wandering across atomic surfaces of single crystal metal and metal oxide surfaces, identifying reactions at specific sites or boundaries of atomic layers [68, 69]. Despite lack of direct evidence for the nature of reaction intermediates in our Pd-catalyzed phosphite oxidation and phosphorylation reactions, enzymatic studies and some observations suggest a possible reaction mechanism, shown in **Figure 3**.

**Figure 3.**
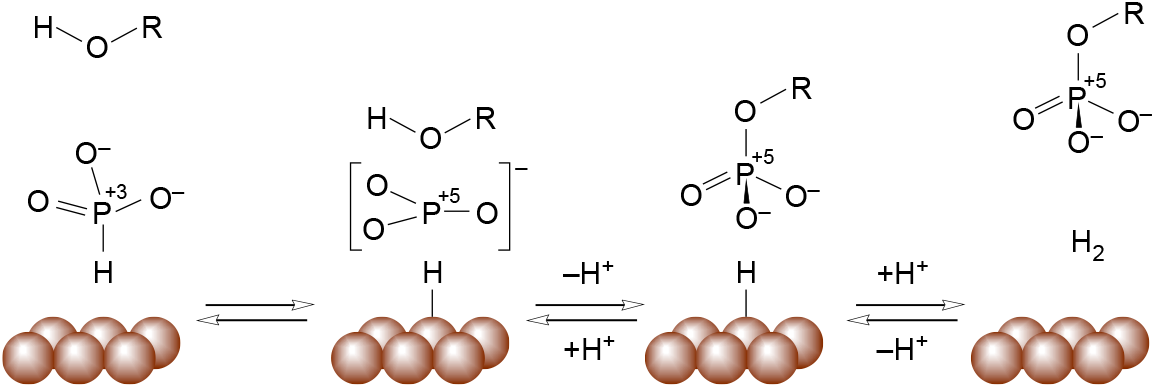
Possible mechanism for phosphite-dependent phosphorylation on metal surfaces. Figure from **Schlikker** *et al*. (2026) [48]. From left to right: Hydride removal generating P^+5^, intermediate formation (metaphosphate, [PO_3_]^−^, shown), P–O bond formation (P–N bond formation in the case of phosphocreatine), formation of phosphate (R=H) or phosphorylated compounds (R=alkyl, phosphoryl, acyl), H_2_ formation.

Justification for the proposed scheme comes from the following. First, we have not observed any cases in which native metals generate phosphorylated products using phosphate as reactant (**Schlikker** *et al*. (2026) [48] and unpublished observations). This indicates that the phosphite oxidation reaction itself is required for phosphorylation, implicating an intermediate of that reaction, rather than its phosphate product, as the phosphorylating agent. Second, enzymatic phosphite oxidation is NAD^+^ dependent in cases studies so far [70, 71] indicating hydride removal in the reaction mechanism [44]. Although neither Ni^0^ nor Pd^0^ can serve as oxidants, they are good hydride acceptors [72–74] and can readily catalyze the synthesis of H_2_ in water via chemisorption in the presence of a strong reductant, which phosphite is, *E*_0_′= –690 mV [75]. Third, metaphosphate is implicated in some enzymatic reactions. Metaphosphate [PO_3_]^−^ is a planar ion with a central P^+5^ atom and is highly unstable in water. **Westheimer** (1981) [76] reports the estimated free energy of the reaction

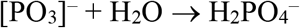

as –107 kJ·mol^−1^, and free metaphosphate is not observed in water [1], hence dissolved metaphosphate can probably be excluded as an intermediate. The dissociative mechanism proposed for *Bacillus* sp. phosphite oxidase [70] involves a metaphosphate intermediate. Metaphosphate is also discussed as an intermediate in enzymatic phosphoester hydrolysis [77]. Fructose-1,6-bisphosphatase has been reported to generate metaphosphate as an enzyme bound reaction intermediate identifiable in the crystal structure [78]. Quantum mechanical studies of phosphoserine phosphatase [79] and mechanistic studies of phosphoglucomutase [80] also suggest the participation of metaphosphate intermediates. Fourth, the accumulation of trimetaphosphate (**Figure 2**) and polyphosphates in our experiments [48] would be compatible with a metaphosphate intermediate, whereby trimetaphosphate (**Figure 2**) would likely have to form on the metal surface;we have no suggestion for metaphosphate-metal interactions. The reaction sketched in **Figure 3** should be reversible, but the equilibrium lies far on the side of phosphate in the presence of water [76].

### Phosphorylation in the early evolutionary process

How does a source of aqueous phosphorylation impact views of bioenergetic evolution [81, 82]? In warm alkaline water characteristic of serpentinizing hydrothermal vents [83], Pd^0^/HPO_3_^2–^ generates the three main kinds of phosphate bonds that occur in core biosynthetic metabolism—the reactions that make the building blocks of life [42, 84]: phosphoesters, phosphoanhydrides and acylphosphates (**Figure 4A**). The occurrence of phosphite in some samples taken from serpentinizing vents [45] and archaean sediments [13] and the occurrence of native metals that can oxidize phosphite (**Figure 2**) suggest that these reactions could have credibly take place in Hadean hydrothermal systems, as outlined in **Figure 4B**. Because serpentinzing systems can have a long life span—the Lost City hydrothermal field is estimated to be ca. 100.000 years old [85] —a geochemical supply of HPO_3_^2–^, in addition to the chemically stable solid state catalysts that phosphorylate organics with it (Ni and Pd), fundamentally change the nature of the “phosphate problem”[7] concerning the origin of metabolism: in essence, it disappears. The ability of native metals to reduce CO_2_ to pyruvate [29], to convert pyruvate to citramalate [28] and to convert pyruvate (plus glyoxylate) into intermediates of the rTCA cycle under hydrothermal vent conditions are well documented [40], these reactions provide 2-oxoacids for reductive aminations [40, 48] and substrates for phosphorylation (**Figure 4B**), as in metabolism. It is now apparent that shiny metals in the same environments can also catalyze the phosphorylation reactions [42, 48] under the same conditions (alkaline, aqueous, ≤100°C), but using phosphite, not phosphate as the phosphodonor.

**Figure 4.**
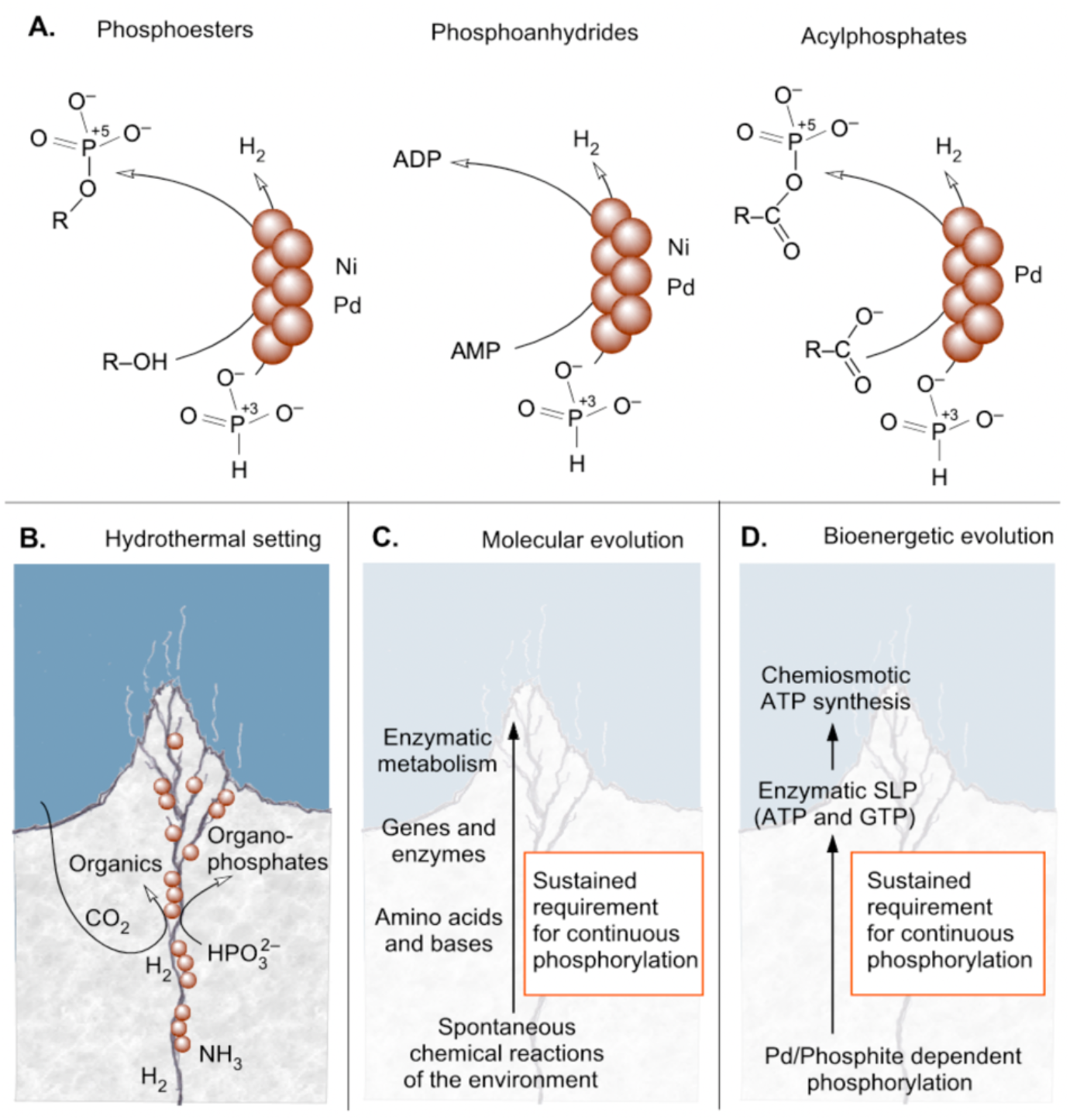
Phosphite-dependent phosphorylation over Pd^0^ in the palaeoecological context of molecular, metabolic and bioenergetic evolution in serpentinizing hydrothermal vents: a chemically reactive site of concomitant organic synthesis and phosphorylation, immune over millennia to disruption by occasional thunderstorms, wind, drought, erosion, UV light, landslides, tsunamis or any combination of the above, and independent of the low solubility of phosphate [90].

This does not, of course, fully bridge the gap to the synthesis of all amino acids, bases, and cofactors, but phosphorylation at the site of substrate synthesis does alleviate two problems inherent to surface environment theories for the ecology of origins. In the cyanosulfidic model [63] various reactions of prebiotic chemistry take place at different times and at very different sites: from impact craters to mountain tops, to streams and pools, drying beds and so forth, with a specific (often nitrile-based, sometimes UV-dependent) chemistry taking place at each site, including phosphorylation. A single, aqueous site for the carbon skeleton synthesis and product phosphorylation in the hydrothermal vent model avoids patchwork-stitchery and product loss inherent to the transport of intermediates from one hypothetical reaction locality to another. In the wet-dry-cycle model for phosphorylation and polymerization [64], products stay in place, as long as neither Hadean thunderstorms nor erosion carries products into the ocean during the thousands to millions of years required to synthesize free-living from the elements. A criticism of hydrothermal vents is that they harbor water, so that products of synthesis would be carried to the ocean, but the concentrating properties of inorganic microcompartments at vents in the presence of temperature gradients (thermophoresis) is well-known [86, 87].

Upon reflection, those issues are actually minor. Thinking through the origins problem one step further, from simple organic synthesis to macromolecular polymer formation, it is evident that whatever environmental mechanism of phosphorylation operated before the origin of genes and enzymes, it had to operate *continuously at the required site of synthesis until enzymes arose that could usurp the function of the environmental phosphorylation process* (**Figure 4C**). This puts severe constraints on the environment and nature of prebiotic phosphorylation, because it means that the environmental phosphorylation mechanism had to be compatible with and coexist with, early generations of proteins (enzymes) until these could themselves provide phosphorylation and it had to operate to support genetically encoded enzyme synthesis on ribosomes. No enzymes are known that harness oven-drying, phosphides, P_2_O_5_ or condensing agents such as cyanoformamid as a source of energy for phosphorylation. Moreover, no enzymes would preserve their activity in such harsh conditions, they would irreversibly denature at the first phosphorylation reaction. By contrast, the enzyme AMP-dependent phosphite dehydrogenase *does* harness the energy contained in the phosphite oxidation reaction to perform substrate level phosphorylation [44] in aqueous solution (bacterial cytosol). In the ADP-forming reaction in **Figure 1**, Pd^0^ merely substitutes for the AdpA enzyme and its NAD^+^ cofactor [44]. Until enzymes invented means to harness environmental energy in such a way as to *replace abiotic phosphorylation*, there existed a non-negotiable demand on the environment to provide a sustained supply of continuous phosphorylation that was compatible with enzymatic chemistry, as indicated in **Figure 4C. Fontecilla**-**Camps** (2019) [88] has emphasized the need for continuity in the environment of early chemical evolution. Aqueous environments and phosphite oxidation afford continuity, harsh chemistry or oven-drying for phosphorylation do not.

One might ask what is the ultimate source of the energy required to synthesize phosphite for early chemical evolution? Two sources seem possible. Either a portion of the phosphorus on the Earth’s early crust following the Moon forming impact was present as phosphite, or (more likely) phosphite was produced from phosphate minerals *in situ* in serpentinizing hydrothermal vents, which have been in existence for as long as liquid water has existed [20] and which generate the extremely reducing conditions needed to convert phosphate to phosphite (*E*_0_′= –690 mV). In the latter case, the source of energy was the same as today: rock water interactions [89] and the amount of phosphite and Pd^0^ required (like the amount of H_2_ required) to fuel early metabolic reactions was governed by the local environment of specific hydrothermal vents (**Figure 4**), not global ocean or atmospheric concentrations.

### A starting point for the bioenergetic process

The new findings impact discussion on the origin of bioenergetics. There are only two sources of net phosphorylation known in life: substrate level phosphorylation and chemiosmotic gradient harnessing via ATP synthases [3, 91]. Substrate level phosphorylation (SLP) involves stoichiometric reactions of organophosphates with sufficient group transfer potential to generate ATP (or ADP from AMP, or GTP from GDP). Such organophosphates are typically acyl phosphates such as 1,3-bisphophoglycerate, succinyl phosphate or acetyl phosphate and the like [55]. SLP is the far simpler and obviously more ancient of the two [92], because the ATP synthase is a large protein, roughly 1/5^th^ the mass of a ribosome, that requires 4 ATP per peptide bond for its own synthesis. It is self-evident that the ATPase cannot have supplied the ATP required for its own origin (and the ribosome that synthesized it) (**Figure 4D**) and a case can be made that GTP, generated via SLP, was the energy currency at the origin of genes, before there were ATP synthases [93]. But for SLP to work, there has to be a supply of organophosphates capable of SLP. Acetyl phosphate has been discussed as a candidate for such a primordial bioenergetic driver [15], mainly because of its essential role in the acetyl CoA pathway of acetogens [91]. Abiotic synthesis of acetyl phosphate has been previously reported [94–96], involving, however, reactants unlikely to have formed by abiotic means. The synthesis of acetyl phosphate reported by **Schlikker** *et al*. (2026) [48] involves acetate, which is readily synthesized from H_2_ and CO_2_ over a variety of solid state metal catalysts under hydrothermal vent conditions [29], phosphite and Pd^0^.

As with metabolic evolution, whatever environmental mechanism of phosphorylation that operated before the origin enzymatic SLP, *it had to operate continuously at the required site of synthesis until enzymes arose that could usurp the function of the environmental phosphorylation process* (**Figure 4D**). And even then, those first enzymes of SPL required an environmentally (abiotically) supplied organophosphate or inorganic phosphodonor capable of phosphorylating ADP. We are suggesting that phosphite over Pd^0^ is that supply. It generates acetyl phosphate at 8 mM, exceeding the ca. 3 mM concentration in *E. coli* cytosol (**Figure 1**) and very close to that of ATP in *E. coli* cytosol, 9.6 mM [56]. A supply at that level could support the origin of genes and proteins.

Phosphite oxidation over Pd^0^ presents a broad-specificity, physiologically sufficient, geologically sustainable and environmentally credible source of phosphorylation at origins. It can also generate acyl phosphates and, once they arise in metabolism, nucleoside phosphates such as ADP, which serve as additional phosphodonors in non-enzymatic phosphorylation reactions [97]. This would generate short, exergonic phosphotransfer cascades at origins, similar to the short phosphotransfer cascades in prokaryotic cells [98]. Phosphite oxidation by shiny metals offers a credible solution to the phosphate problem in that it involves reactants and catalysts generated under the continuously stable, dark and reducing conditions of serpentinizing hydrothermal vents, where the organic substrates for phosphorylation are themselves produced. For those who have argued for decades in favor of the evolutionary significance of hydrothermal vents in spawning life [99–101], phosphite over Pd^0^ presents progress on the origins problem, as it unites CO_2_ fixation, phosphorus based bioenergetics, microbial metabolism and continuously favourable thermodynamics in a single physiologically relevant environment, where anaerobic autotrophs still thrive [83, 102] on nutrients supplied by serpentinization.

## Acknowledgements

We thank Maximilian Burmeister and Eleni Dafni for help in the laboratory.

