## Supplemental Information for "Metal-catalyzed phosphorylation by phosphite at the origin of bioenergetics"

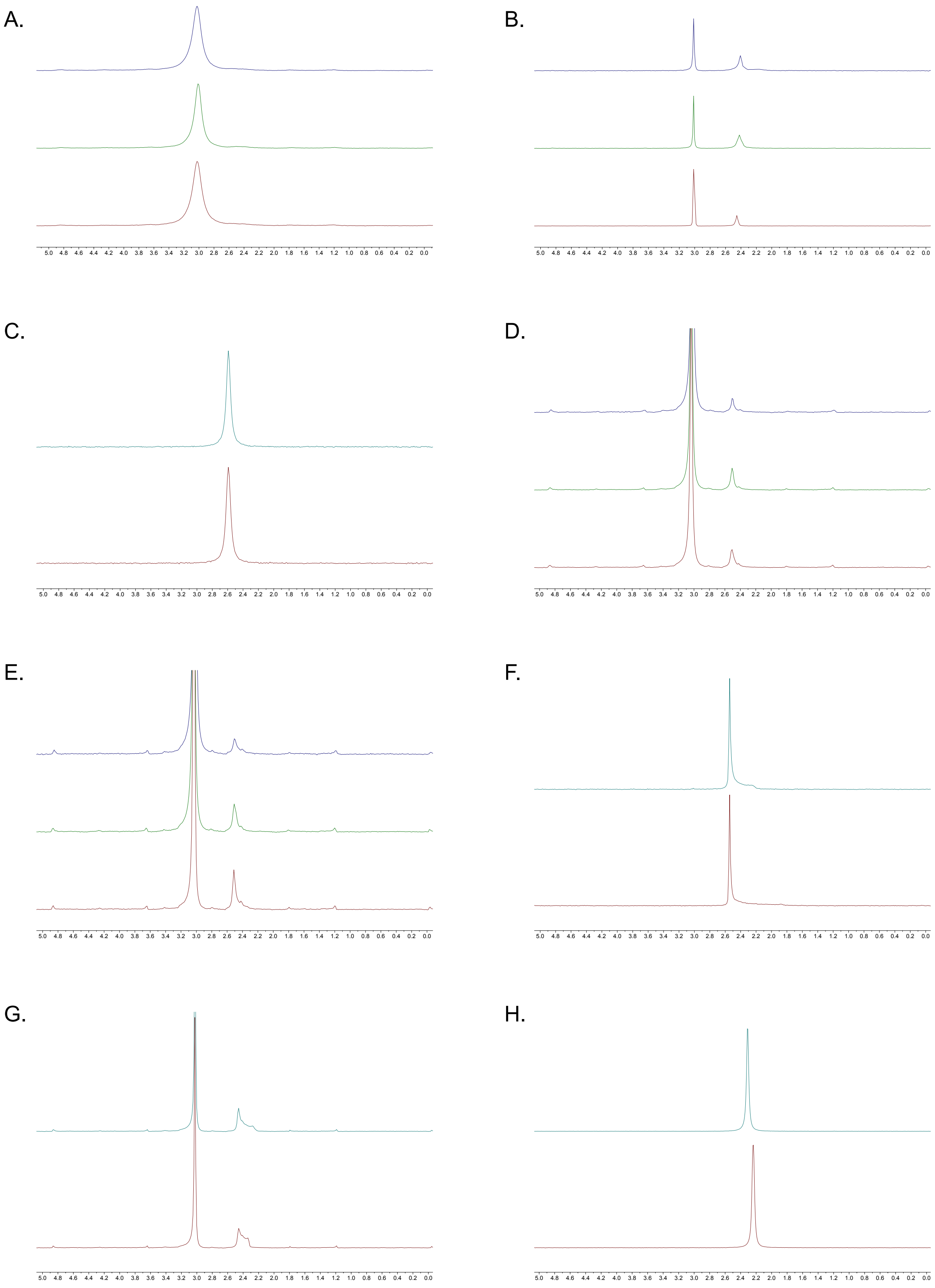
A.
B.
C.
D.
E.
F.
G.
H.

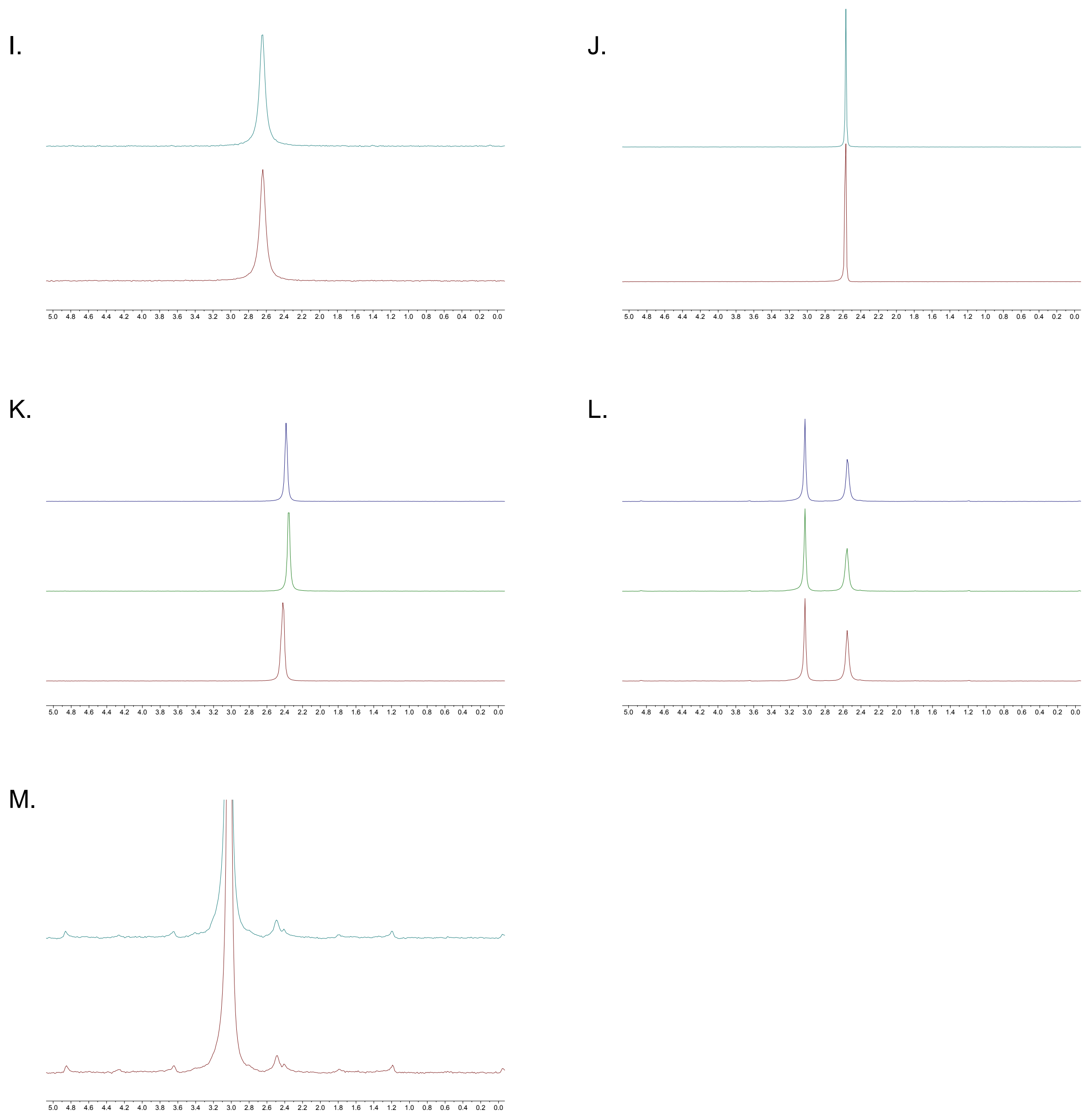


**Figure S1.** $^{31}$P-NMR spectra corresponding to the phosphite oxidation experiments shown in **Figure 2A** after incubation at 100 °C for 72 h with different metal catalysts. All reactions contained 200 mM phosphite and 0.15 mmol catalyst, except for Pd/C, where 1.5 mmol catalyst was used. Spectra are shown for (**A**) control without catalyst, (**B**) $Pt^0$, (**C**) $Pt^0/Al_2O_3$, (**D**) $Ir^0$, (**E**) $Rh^0$, (**F**) $Rh^0/Al_2O_3$, (**G**) $Ru^0$, (**H**) $Ru^0/C$, (**I**) $Ru^0/Al_2O_3$, (**J**) $Pd^0$, (**K**) $Pd^0/C$, (**L**) $Ni^0$, and (**M**) $Ni^0/C$.

A.

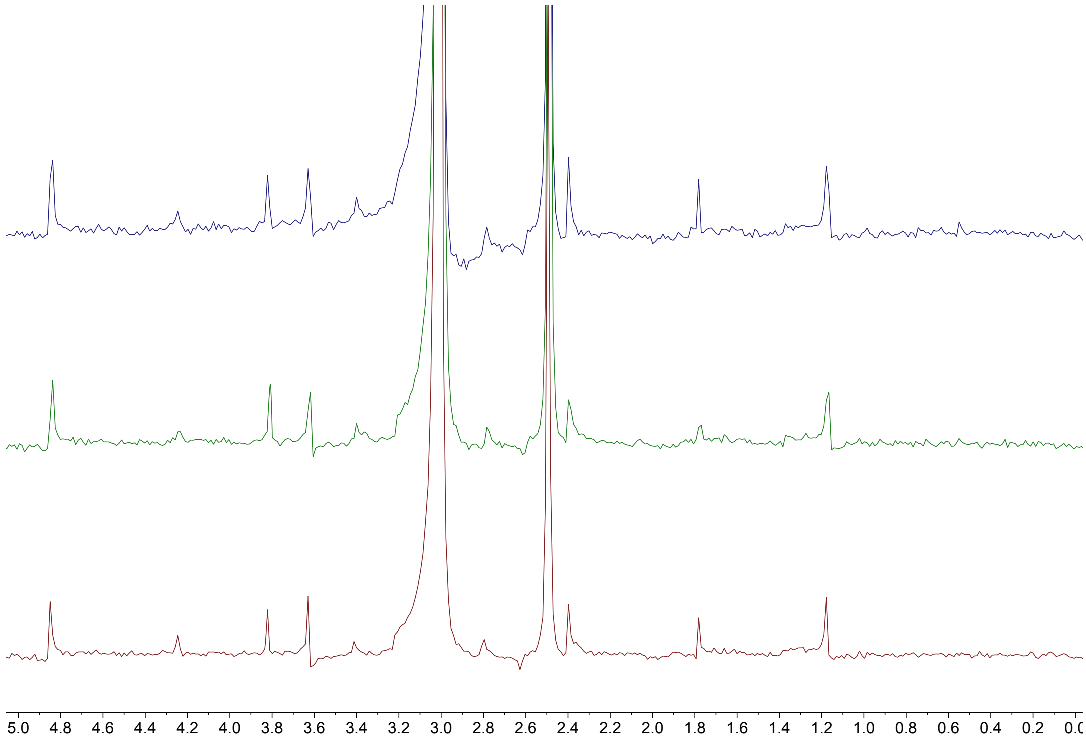


B.

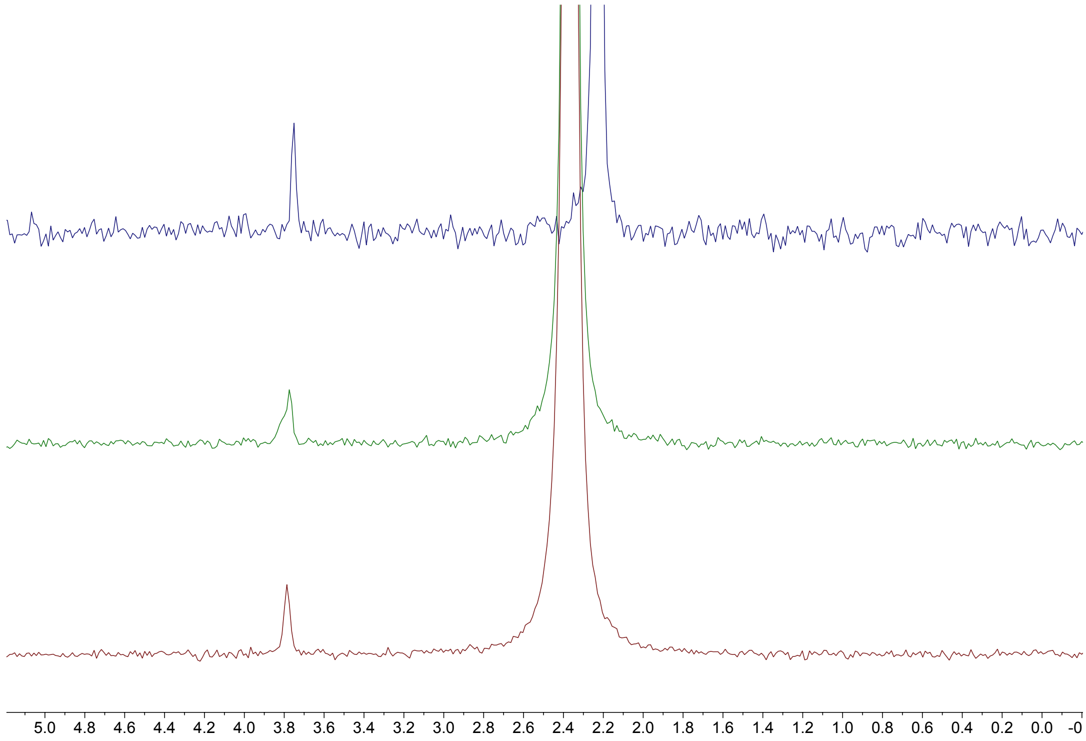


**Figure S2.** Raw $^{31}P$-NMR spectra corresponding to the phosphoserine formation experiments shown in **Figure 2B** after incubation at 50 °C for 18 h. All reactions contained 100 mM serine and 200 mM phosphite. **A**, shows the reaction with $Pd^0$, and **B**, the reaction with Pd/C.

A.

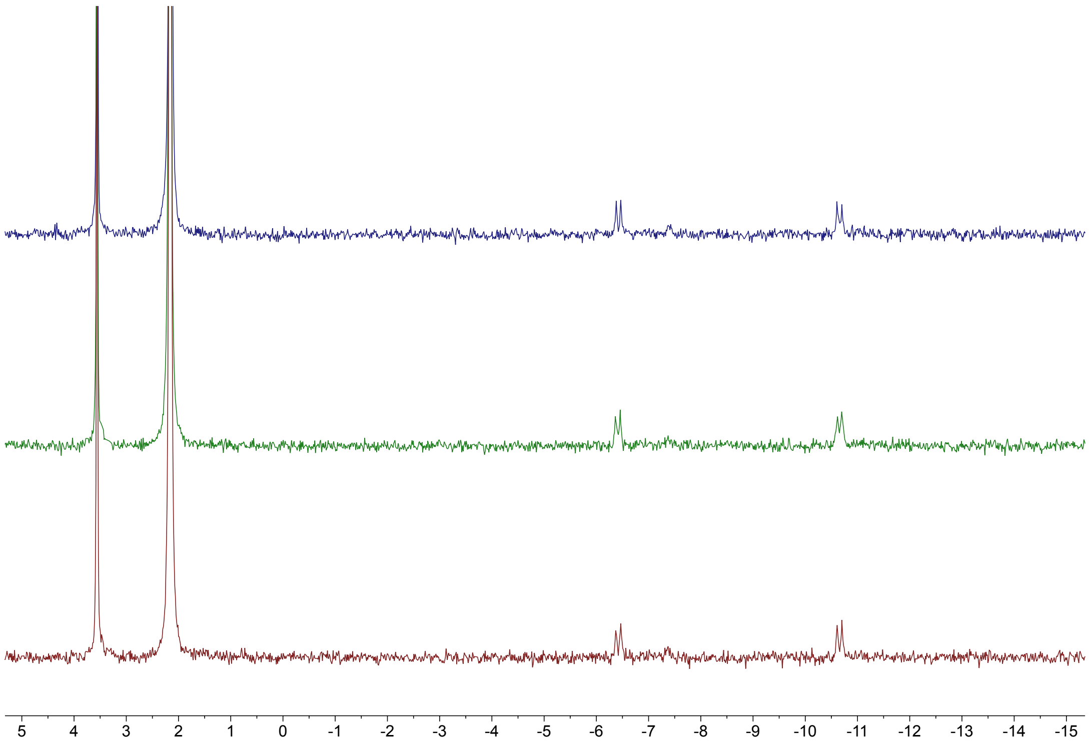


**Figure S3.** Raw $^{31}$P-NMR spectrum corresponding to the AMP phosphorylation experiment shown in **Figure 2C** after incubation at 50 °C for 18 h. The reaction contained 100 mM AMP and 200 mM phosphite.
